# Primordial endocrine cells uncouple sexual differentiation from sex-chromosomes

**DOI:** 10.64898/2026.09.21.753345

**Authors:** Tom Levy, Melody Wahl, Uzi Hadad, Benyamin Rosental, Rivka Manor, Simi Weil, Eliahu D. Aflalo, Chiara Benvenuto, Amir Sagi

**Author notes:** Corresponding Authors E-mail: Tom Levy, Amir Sagi.

## Abstract

Sexuality among the animal kingdom is remarkably plastic, spanning from gonochorism to sequential or simultaneous hermaphroditism. Over 70 years ago, Charniaux-Cotton hypothesized that sexual plasticity in crustaceans is mediated by primordial cells present in both sexes, which differentiate into an androgenic gland (AG) only in males. Although AG engraftments can masculinize genetic females, it remained unclear whether this reflects long-term integration of donor cells within the host or activation of host primordial AG progenitors. Using the freshwater prawn as an exceptional case of sexual plasticity, where AG-cell transfer yields functional neo-males bearing only female sex-chromosomes, we show, through cellular-resolution approaches, that the AG arises from primordial cells present in every individual, indicating autosomal machinery of sexual differentiation completely independent from the sex-chromosomes.

## Introduction

Sexual plasticity, the capacity of individuals to express alternative sexual phenotypes, is widespread across the animal kingdom and spans a spectrum from strict gonochorism through full hermaphroditism to asexual parthenogenesis (Bachtrog et al., 2014; Benvenuto & Weeks, 2020; Scholtz et al., 2003; Todd et al., 2019). Across animals, this plasticity is manifested in diverse contexts, from environmentally induced sex determination (Bachtrog et al., 2014; Estermann et al., 2020; Flament, 2016), to naturally occurring sex change (Goikoetxea et al., 2021; Pla et al., 2022; Todd et al., 2019) and experimentally induced endocrine sex reversal (Flament, 2016; Goikoetxea et al., 2021).

In crustaceans, sexual plasticity is governed by the androgenic gland (AG), the master endocrine regulator of male differentiation. Classical studies by Charniaux-Cotton (1954, 1955, 1957) demonstrated that AG removal feminized males, whereas AG implantation into immature females induced complete masculinization. Gonadal transplants alone had no such effect, establishing the AG as the exclusive source of an endocrine component responsible for male sexual differentiation in crustaceans (Charniaux-Cotton, 1953, 1954, 1955). This conserved role was subsequently confirmed across decapod species, including prawns, crayfish, shrimp and crabs (Alfaro-Montoya et al., 2016; Barki et al., 2003; Cui et al., 2005; Kato et al., 2015; Khalaila et al., 2001; Manor et al., 2004; Nagamine et al., 1980a, 1980b; Taketomi & Nishikawa, 1996).

In decapod crustaceans the AG acts as a sexual switch (Levy et al., 2020; Miao et al., 2023; Rosen et al., 2010; Sagi et al., 1997; Tan et al., 2020; Villanueva et al., 2022) through the insulin-like androgenic gland hormone (IAG) (Manor et al., 2007), termed the IAG-switch (Levy & Sagi, 2020). The ON/OFF state of the switch dictates the sexual differentiation trajectory, ultimately leading to distinct sexual maturation: ON directs male development (Levy et al., 2016), whereas OFF permits female development (Miao et al., 2023). Experimental or natural manipulations of the IAG-switch in decapods has established its role in directing sexual differentiation path (Chen et al., 2022; Khalaila et al., 2001; Levy et al., 2020; Levy et al., 2021; Miao et al., 2023; Rosen et al., 2010; Rubiliani & Payen, 1979; Veillet & Graf, 1958). While partial sex-manipulations occurred in many decapods following the manipulation of the switch [reviewed in Levy and Sagi (2020)], the major species in which a completely functional sex-reversal was successful is the giant freshwater prawn *Macrobrachium rosenbergii*. In *M. rosenbergii*, which follows a WZ-ZZ sex determination system (females are WZ and males are ZZ; Fig. 1) (Malecha et al., 1992; Sagi & Cohen, 1990; T Ventura et al., 2011), manipulation of the switch has enabled remarkable demonstrations of sexual plasticity (Levy et al., 2016; Levy et al., 2019; Ventura et al., 2012). A major step in detecting successful IAG-switch manipulations was the development of genetic sex markers (T Ventura et al., 2011), which enabled W/Z genotyping and confirmed the sex reversal of genetic ZZ males into functional ZZ neo-females following RNAi-mediated *IAG* silencing (Ventura et al., 2012). Conversely, a single injection of suspended hypertrophied AG cells at an early post-larval stage induced complete functional sex reversal of genetic WZ females into fully reproductive WZ neo-males, which were subsequently crossed with normal WZ females to generate WW females (Levy et al., 2016). Further manipulation through injection of suspended AG cells into genetic WW females produced fully functional WW neo-males (Levy et al., 2019). Together, these manipulations showed that all possible genotypes (ZZ, WZ, and WW) can manifest phenotypically as either sex (Fig. 1A).

**Fig. 1.**
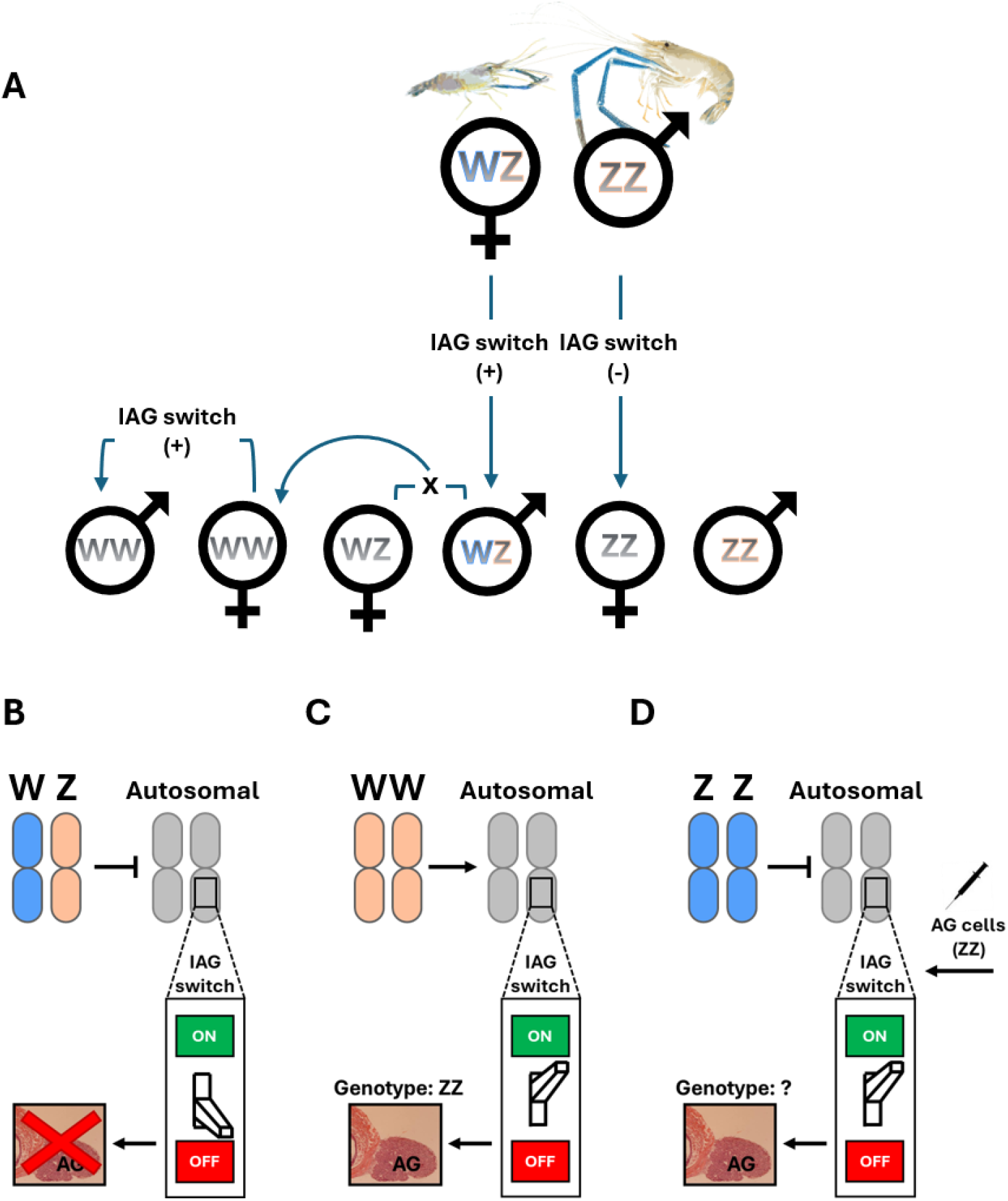
Sexual plasticity in prawns. (A) Manipulation of the IAG-switch enables the production of male and female phenotypes from each genetic sex genotype (WZ, ZZ, WW). In unmanipulated animals, the IAG-switch is OFF in WZ females and ON in ZZ males. Activation of the switch in WZ females produces WZ neo-males (Levy et al., 2016), whereas suppression of the switch in ZZ males produces ZZ neo-females (Ventura et al., 2012). Crosses between WZ neo-males and normal WZ females yield WW offspring, including WW females (Levy et al., 2016). Subsequent activation of the switch in WW females produces fully functional WW neo-males (Levy et al., 2019). Plus (+) and minus (-) signs indicate IAG-switch activation and suppression, respectively. (B-D) The putative interaction between the sex-chromosomes and the IAG-switch which is predicted to reside within autosomal chromosomes. (B) The pathway from a WZ genotype to the suppression of the AG development and (C) the pathway from a ZZ genotype to the development of an AG. (D) The procedure using ZZ AG cell transplantation to bypass the AG suppression and trigger AG development whose genotype is unknown in a WW animal.

The emergence of a functional AG in WW neo-males, individuals entirely lacking the Z chromosome, suggests a unique opportunity to investigate a fundamental question regarding the origin of the gland (Fig. 1D): whether sex reversal following AG cell transfer (from ZZ donor) is driven by long-term integration of donor cells in the host, or by a transient inductive signal that triggers endogenous host primordial cells to develop into a functional AG. More broadly, this question bears on the cellular basis of sexual plasticity: whether all individuals, regardless of sex-chromosome constitution, retain, at early stage of development, primordial cells competent to generate the male endocrine system.

This echoes a deeper hypothesis rooted in Charniaux-Cotton’s original observations, that sexual phenotype is determined by the developmental fate of primordial cells with the capacity to form an AG de novo rather than by sexual genotype per se (Charniaux-Cotton, 1959; Charniaux-Cotton, 1962). Despite decades of transplantation experiments, this hypothesis has not been tested. Here we address this question directly: our ability to generate a unique WW neo-male system, together with genomic sex markers for determining sex-chromosome composition and high-resolution AG-cell isolation by laser capture microdissection (LCM) and fluorescence-activated cell sorting (FACS), enabled direct testing of this hypothesis, revealing the cellular origin of the gland and providing insight into the cellular foundations of sexual plasticity.

## Results

### Active AG in WW neo-males is comparable to ZZ males

Histological sections of the base region of the fifth pereiopods from WW and ZZ males revealed fully developed AGs in both. Immunohistochemistry using Mr-IAG specific antibodies confirmed active Mr-IAG hormone production in both genotypes (Fig. 2), indicating that AG activity in WW neo-males is comparable to wild-type ZZ males despite the absence of the masculine Z chromosome.

**Fig. 2.**
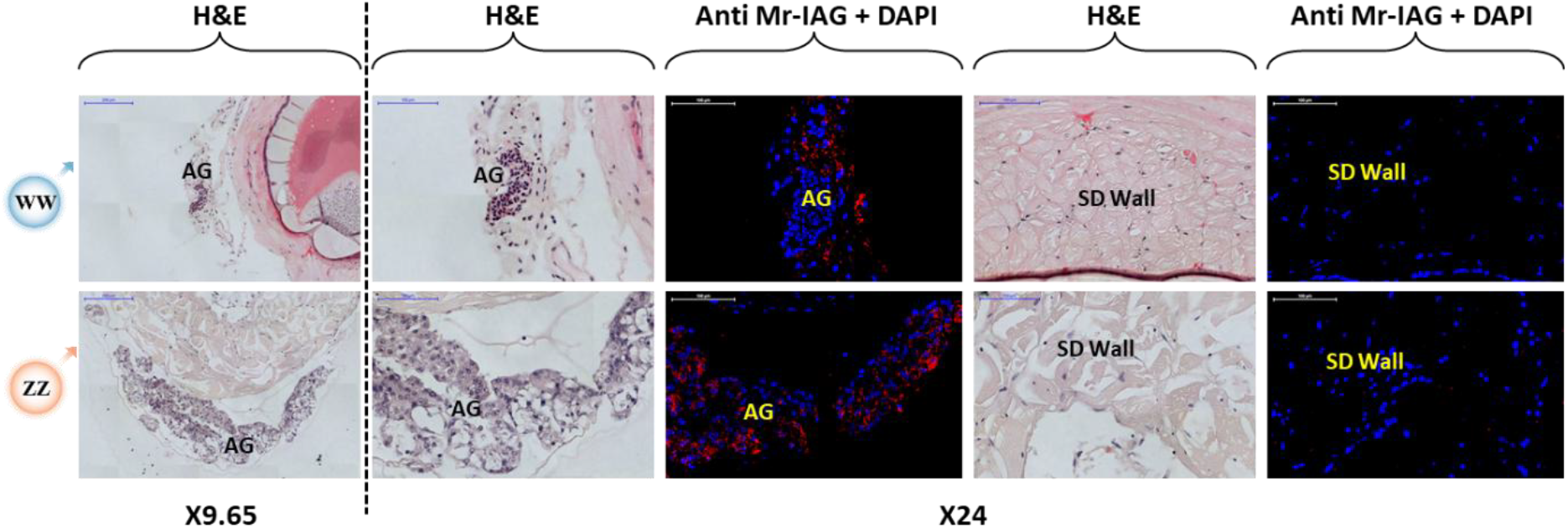
Histological sections and immunohistochemistry of fifth pereiopods from WW (top) and ZZ (bottom) males. The androgenic gland (AG) and sperm duct (SD) wall are marked. Histological sections were stained with hematoxylin and eosin (H&E). Immunohistochemical sections were stained with Alexa 546-conugated anti-Mr-IAG antibodies (red) and with DAPI (blue) for nuclei counter-staining. Scale bars and magnifications (200 µm, ×9.65; or 100 µm, ×24) are indicated in each image.

### Cellular *Mr-IAG* expression pattern is unaffected by sex-chromosomes

Cell sorting resolved four distinct AG cell types from both ZZ and WW males based on size and granularity (Fig. 3A, 3B, 3E). Population 1 contained large cells (20 µm) and populations 2, 3 and 4 contained smaller cells (5-10 µm). *Mr-IAG* expression was similar across populations 1, 2 and 4, while population 3 showed significantly reduced expression, a pattern identical in both ZZ and WW males (Fig. 3C and 3D). This demonstrates that AG cellular composition and IAG expression might vary among different AG cell types but are unaffected by the genotypic composition of the sex-chromosomes.

**Fig. 3.**
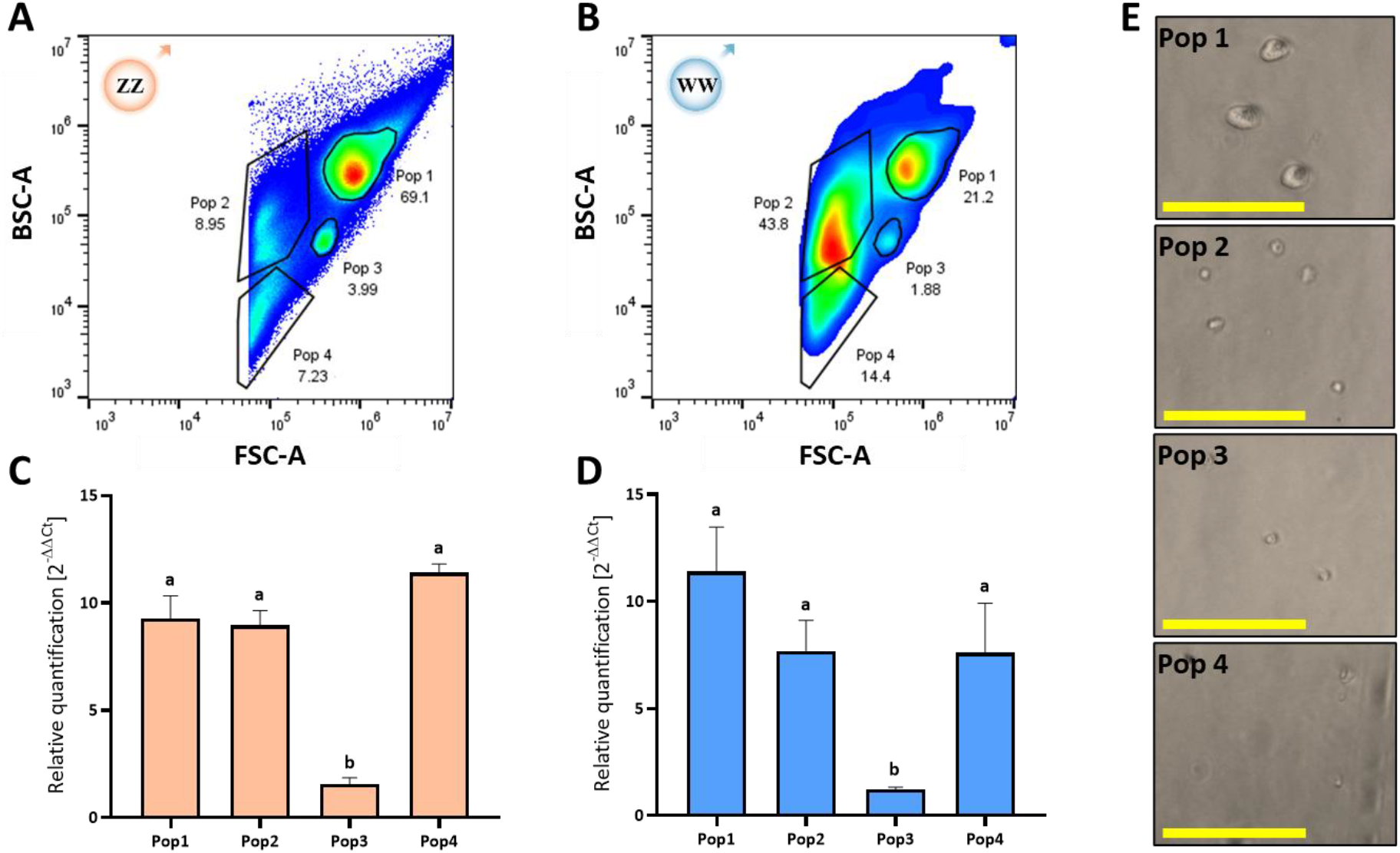
*Mr-IAG* expression in sorted AG cell types. FACS plot showing the main four cell populations that were sorted from (A) ZZ males and (B) WW neo-males. (C) Relative *Mr-IAG* quantification, using qPCR, in the four sorted cell populations showed similar pattern in (C) ZZ males and (D) WW neo-males. Statistical significance (*P* ≤ 0.05) for each plot is indicated with letters. (E) Representative photos under light microscopy show morphological differences between the four sorted AG cell populations (bars=100 µm).

### Neo-male AG originates from host WW primordial cells

Laser-capture microdissection of AG cells from histological sections from ZZ and WW males showed that their AG cells bear ZZ and WW genotypes, respectively, matching the genotype of the testes and muscle tissues from the same individuals (ZZ in ZZ males and WW in WW males; Fig. 4). This confirms that following transient IAG induction by donor AG cell suspension transfer, the AG develops in the host from endogenous primordial cells, regardless of sex-chromosome genotype.

**Fig. 4.**
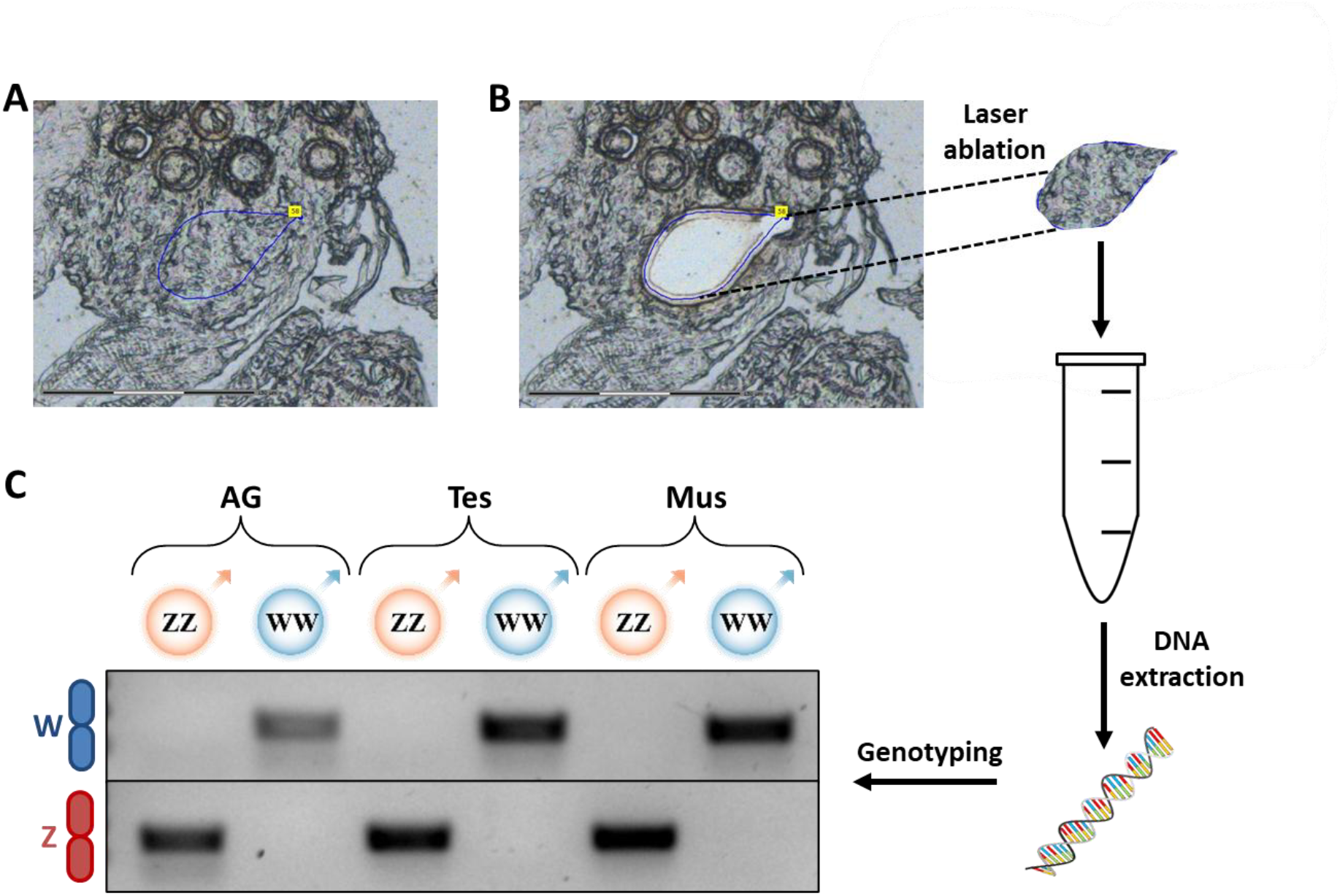
Isolation and genotyping of the AG cells in ZZ males and WW neo-males. Representative photo of AG histological section (A) before and (B) after laser ablation (bars=150 µm). DNA was extracted from the laser-ablated fragments containing the cells. (C) PCR amplification of sex-specific genomic markers (W – top and Z – bottom) using DNA from laser-ablated AG cells as a template. DNA isolated from testis cells (Tes) and pleopod muscle (Mus) of the same WW and ZZ males served as controls.

## Discussion

Our results show that unlike intact tissue grafts, suspended donor cells are unlikely to form a donor-derived gland. The AG developed in neo-male did not arise through persistent integration of transplanted donor cells but developed de novo from host cells following transient AG-cell induction. This conclusion was enabled using suspended AG cell transfer from ZZ donors into WW females to generate WW neo-males (Levy et al., 2016; Levy et al., 2019), and high-resolution AG cell isolation and genotyping. The WW genotype of the AG in WW neo-males, matching host gonad and muscle tissues, demonstrates that the transferred donor AG cells act as an inductive trigger rather than as the structural source of the gland, suggesting that the endocrine signal delivered by the transplanted cells is sufficient to induce host AG formation.

The nature of the inductive signal delivered by transplanted AG cells remains to be directly established, but the endocrine architecture of the system constrains the plausible mechanisms. IAG acts through membrane-bound tyrosine kinase insulin receptors on target tissues (Aizen et al., 2016; Guo et al., 2018; Sharabi et al., 2016), and a signal of this kind is necessarily humoral rather than cell-autonomous: it cannot enter recipient cells directly to act as a transcriptional regulator. In the isopod *Armadillidium vulgare*, androgenic hormone expression precedes visible AG primordium differentiation and has been proposed to act in an autocrine manner, driving AG formation during a discrete developmental window. However, disruption of insulin signaling by *Wolbachia*, likely through suppression of upstream regulating neurohormones rather than direct receptor inactivation, prevents this autocrine loop and results in feminization (Herran et al., 2020; Herran et al., 2021). These observations suggest that IAG itself, or a related factor secreted by mature AG cells, may constitute the inductive signal that recruits host primordial cells, consistent with the model we suggest here for decapods. According to this model, the transplanted AG cells provide a transient local source of IAG, and potentially other factors, that trigger autocrine or paracrine amplification in host progenitors during the competence window.

A transient inductive signal acting on competent progenitors within a restricted developmental window is a recurring logic in vertebrate endocrine organogenesis. Progenitor cells pass through sequential, temporally distinct states with differing lineage biases, and signals commit cells to an endocrine fate; prolonged or mistimed exposure disrupts correct differentiation (Bastidas-Ponce et al., 2020; Rebourcet et al., 2014; Scavuzzo et al., 2018). The crustacean IAG-switch parallels this logic: a secreted peptide from a differentiated endocrine source is suggested to act during a defined early window to instruct competent progenitors toward an alternative cellular fate, in this case the male endocrine identity, rather than supplying that identity directly.

This study provides direct cellular evidence for Charniaux-Cotton’s longstanding hypothesis that crustacean individuals, regardless of sex-chromosome composition, harbor primordial cells competent to form a functional AG (Charniaux-Cotton, 1959; Charniaux-Cotton, 1962). In females, these cells may normally remain inactive or be diverted from the AG pathway, whereas exposure to the appropriate inductive endocrine context during an early developmental window can trigger their differentiation into a functional secretory gland. Sexual plasticity in this system therefore reflects not only phenotypic reversibility, but also the persistence of AG-competent primordial cells in every individual, indicating that cellular potential for alternative sexual differentiation is retained across genotypes, for a critical time window. The fate of these cells is governed by the IAG-switch ON/OFF state. Hormonal context, rather than genomic sex, determines whether they differentiate into a secretory gland or remain quiescent. Moreover, the fact that WW males, entirely lacking the Z chromosome, develop fully functional AGs with IAG expression patterns comparable to ZZ males, supports a model in which sex-chromosomes direct sexual fate upstream, whereas the downstream cellular and endocrine machinery required to drive sexual differentiation is shared across genotypes and likely resides predominantly within the autosomal system. This framework is consistent with hermaphrodite animals, which can develop male and female gonads, simultaneously or sequentially, without chromosomal sex determination and especially supported in decapods by the exclusive occurrence of protandric and simultaneous hermaphroditism (Baeza, 2018; Benvenuto & Weeks, 2020; Bortolini & Bauer, 2016; Levy et al., 2020).

Unlike mammals, in which endocrine cell lineages involved in sexual differentiation are embedded within the gonad (Estermann et al., 2020; Mäkelä et al., 2019; Rotgers et al., 2018), the crustacean AG is an extragonadal endocrine organ (Charniaux-Cotton, 1962). This anatomical separation between the endocrine switch and the gametogenic tissue allows manipulation of sexual differentiation independently of the gonad, making it possible to generate genotype–phenotype combinations absent from nature, including functional males lacking male sex-chromosomes (Levy et al., 2019) and functional females lacking female sex-chromosomes (Ventura et al., 2012). It also opens a broader view of sexual plasticity in which chromosomal sex may direct the developmental trajectory, but alternative sexual phenotypes can emerge from shared cellular substrates when the endocrine context is redirected (Abberbock et al., 2026; Estermann et al., 2020; Mäkelä et al., 2019; Rotgers et al., 2018; Zarkower, 2006).

This study places sexual plasticity at the level of endocrine organogenesis. The alternative sexual phenotype is not achieved only by modifying an existing hormonal pathway, but by inducing de novo formation of the male endocrine organ from host primordial cells. Thus, the crustacean IAG-switch provides a tractable model for understanding how chromosomal sex, cellular competence, and endocrine induction interact to generate phenotypic sexual outputs and plasticity.

## Acknowledgements

We thank Enzootic Ltd for supplying prawns for this research.

## Author contributions

T.L. and A.S. conceived the project and designed the overall study. M.W. and T.L. performed the statistical analyses. U.H. and B.R. provided access to, guidance on, and technical support for laser-capture microdissection, flow cytometry, and microscopy. S.W. and E.D.A. performed tissue dissections. T.L. and E.D.A. performed histological sectioning and immunohistochemistry. T.L., M.W., and R.M. conducted molecular experiments, including cell sorting, DNA and RNA extractions, PCR, and qPCR. C.B. contributed to writing and provided expertise on hermaphrodite species. T.L., M.W., and A.S. drafted the manuscript, with all authors contributing to the review and editing process and providing final approval for publication.

## Materials and Methods

### Animal husbandry and sex reversal of WW females into WW neo-males

*M. rosenbergii* males were reared in 600-L tanks at 28 ± 2 °C with constant aeration, a light regime of 14:10 (L/D) and were fed ad libitum (shrimp pellets comprising 30 % protein) at the aquaculture facility at Ben-Gurion University of the Negev.

Sex reversal of WW females into WW neo-males was performed as previously described (Levy et al., 2016; Levy et al., 2019). Briefly, the male donors were endocrinologically manipulated, leading to AG hypertrophy after 10 days, followed by AG dissection and production of AG cell suspension. These AG cells were transplanted (using a micro-injector under a light microscope), in an amount of ∼3,000 cells per female prawn, into the abdomens of WW females, at an age of less than 30 days post larvication. The injected prawns were reared in 600-L tanks for grow-out.

### Histology and immunohistochemistry

Testes, together with the proximal sperm duct and fifth pereiopods (the location of the AG), were dissected from WW neo-males and ZZ males (∼1 year of age). Tissue samples were fixed in 4 % buffered formalin for 48 h in room temperature. Samples were then gradually dehydrated through a series of increasing alcohol concentrations, incubated with xylene and embedded in Paraplast (Kendall, Mansfield, MA) according to conventional procedures. Five-micrometer-thick sections were cut and laid onto silane-coated slides (Menzel-Gläser, Braunschweig, Germany). For morphological observations, one of five consecutive slides were hematoxylin and eosin-stained as previously described (Levy et al., 2016) while other selected slides were analyzed by immunohistochemistry using Alexa 546-conugated anti-*Mr-IAG* antibodies, as well as DAPI for nuclear counter-staining, as previously described (Levy et al., 2016; Tomer Ventura et al., 2011).

### Cell sorting and in-vitro expression of *Mr-IAG*

Hypertrophy of AGs from sexually-mature *M. rosenbergii* WW neo-males (n=4) and ZZ males (n=5) was achieved by surgical removal of the neuroendocrine X organ-sinus gland complex, located in the eyestalk. Eight days post eyestalk ablation, the animals, weighing 87.3 ± 55 g, were anesthetized for 10 min in ice-cold water supplemented with 0.2 % hypochlorite for disinfection purposes and their AGs were dissected under light microscope. AG cell suspension was produced as previously described (Levy et al., 2016). Briefly, AG cells were separated using enzymatic dissociation, followed by centrifugation and washing steps (3 times) and re-suspended in 1 ml of medium [Leibovitz L-15 medium with L-glutamine, 10 % (v/v) fetal bovine serum, and penicillin-streptomycin solution]. The re-suspended cells were placed on ice until FACS analysis. Cell sorting was carried out using Sony MA900 FACS instrument based on size (forward-scatter; FSC) and granularity (back-scatter; BSC). Flow cytometry data were analyzed using Flowjo v10.10 Software (BD Life Sciences). A total of 100,000 cells from each population were sorted into 750 µL of TRI-Reagent LS (Sigma-Aldrich) and RNA extraction was carried out according to the manufacturer instructions.

Total RNA was extracted with the TRI Reagent® LS RNA isolation kit (Sigma-Aldrich, Saint Louis, MO, USA), according to the manufacturer’s instructions. RNA was extracted from sorted AG cell populations of WW neo-males (3 replicates per cell population) and ZZ males (4 replicates per cell population), with each replicate representing an independent biological sample. Complementary DNA (cDNA) was synthesized using a qScript cDNA Synthesis Kit (Quantabio, MA, USA) with 100 ng of extracted total RNA, according to the manufacturer’s instructions.

Relative quantification of *Mr-IAG* transcript levels was performed by quantitative PCR (qPCR) using gene-specific primers for *Mr-IAG* (forward: 5′-GCCTTGCAGTCATCCTTGA-3′; reverse: 5′-AGGCCGGAGAGAAGAATGTT-3′) and *Mr-18S* (forward: 5′-CACCTCTCGCGTCGTAGTAC-3′; reverse: 5′-TGCCCGAATGTTACCTGCAT-3′). Universal qPCR BIO Probe Blue Mix (PCR Biosystems, London, UK) and designated qPCR probes (Biolegio) were used for *Mr-IAG* (5′-TTCCCTCTTCCTTATATTTCG-3′) and *Mr-18S* (5′-CACCTCTCGCGTCGTAGTAC-3′). *Mr-18S* (GenBank accession no. GQ131934.1) served as the reference gene. The qPCR reactions were performed on a QuantStudio 1 Real-Time PCR System (Applied Biosystems, Foster City, CA, USA).

### Statistical Analysis

*Mr-IAG* relative transcript levels between sorted AG cell populations within each genotype were compared using one-way ANOVA, followed by post hoc Tukey’s HSD test. To meet the assumptions of one-way ANOVA, residuals’ normality was tested using the Shapiro-Wilk test, and homogeneity of variances was tested using Levene’s test. The data were logarithmically transformed to meet the assumptions of proper statistical analysis. All statistical analyses were performed using Statistica v13.5 software (StatSoft Ltd., Tulsa, OK, USA).

### Genotyping of AG cells, testes and muscle tissues

Laser capture microdissection, Zeiss Palm MicroBeam instrument, was used to precisely isolate AG and testes cells from unstained histological sections to avoid contamination from surrounding tissues. Muscle tissues were sampled from the second pleopod of representative animals. Genomic DNA was extracted using REDExtract-N-Amp Tissue PCR Kit (Sigma Aldrich) according to the manufacturer’s instructions, and the genotype of each animal was determined using W/Z-specific genomic markers for *M. rosenbergii*, as previously described (Levy et al., 2016; T Ventura et al., 2011). PCR products were separated on a 2 % agarose gel, stained with SYBR Safe DNA Gel Stain.

